# Metabolic turnover and dynamics of modified ribonucleosides by ^13^C labeling

**DOI:** 10.1101/2021.07.20.453046

**Authors:** Paulo A. Gameiro, Vesela Encheva, Mariana Silva Dos Santos, James I MacRae, Jernej Ule

## Abstract

Tandem mass spectrometry (MS/MS) is an accurate tool to assess modified ribonucleosides and their dynamics in mammalian cells. Yet, MS/MS quantification of lowly abundant modifications in non-ribosomal RNAs is unreliable, and the dynamic features of various modifications poorly understood. We developed a ^13^C labeling approach, ^13^C-dynamods, to quantify the turnover of base modifications in newly transcribed RNA. This turnover-based approach helped to resolve mRNA from ncRNA modifications in purified RNA or free ribonucleosides, and showed the distinct kinetics of *N*6-methyladenosine (m^6^_2_A) versus 7-methylguanosine (m^7^G) in polyA+-purified RNA. We uncovered that *N*6,*N*6-dimethyladenosine (m^6^_2_A) exhibits a distinct turnover in small RNAs and free ribonucleosides when compared to the known m^6^ A-modified large rRNAs. Finally, combined measurements of turnover and abundance informed on the transcriptional versus posttranscriptional sensitivity of modified ncRNAs and mRNAs, respectively, to stress conditions. Thus, ^13^C-dynamods enables studies of origin of modified RNAs at steady-state and their dynamics under non-stationary conditions.

## INTRODUCTION

RNA methylation modulates crucial RNA-protein interactions at various stages of RNA metabolism. Comprehensive studies of individual RNA modifications have been mostly advanced by applications of next-generation sequencing (NGS), which rely on chemical, enzymatic and/or antibody-based detection of modified ribonucleosides (1–3). These methods have provided a wealth of information on the sites of modification across the transcriptome (1–3), which have uncovered the dynamic behavior of *N*6-methyladenosine (m^6^A) mRNA methylation in development (4–8). Notably, dysregulation of m^6^A levels has been recently linked to cancer, ageing and neurodegeneration (9–15), which can occur via metabolic inhibition of m^6^A demethylation (9, 10). Furthermore, various tRNA modifications have been reported to be post-transcriptional sensitive to cellular stress (16–19). A current caveat of NGS-based profiling is the lack of high specificity reagents for every modification (1–3). Also, the effects of transcription, which often predominate in differential expression analyses (20), make it challenging to quantify general changes in methylation levels, with limited insight being obtained into biological associations between various modifications. As a consequence, the dynamic behavior of RNA methylation often remains poorly understood, and complementary approaches are needed to quantify multiple modifications across biological contexts and to assess their associations to transcriptional versus posttranscriptional events.

Tandem mass spectrometry (LC-MS/MS) is a highly accurate tool for analysis of modified RNAs, which has so far been applied primarily in two approaches to identify and quantify RNA modifications (21). The first approach employs LC-MS/MS of intact RNA oligonucleotides to detect multiple modifications with positional information in specific RNA sequences. This approach is chromatographically challenging (22) and requires advanced data mining for unambiguous identification of RNA fragments, and it has so far been applied for comprehensive characterisation of RNA modifications in abundant or short RNAs (22), such as rRNA (23–26), tRNAs (27, 28) and miRNAs (29). The second approach employs LC-MS/MS of ribonucleosides for sensitive quantification of RNA modifications (21). This approach, often combined with stable isotopes, can simultaneously quantify the abundance of multiple modifications in specific RNA classes of interest, being highly suitable to assess the presence of tRNA and rRNA modifications and their dynamics under various biological scenarios (19, 27, 30–35). These methods have so far been applied mainly to study these abundant RNA species because MS detection of mRNA modifications is unreliable due to the heavily modified ncRNAs, which invariably contaminate mRNA pools purified using either polyA+-enrichment or rRNA depletion (36, 37). Therefore, MS approaches are needed that are capable to account for and quantitatively resolve the origin of multiple RNA modifications in purified RNA. Moreover, the dynamic behavior of RNA modifications is insufficiently explained solely by changes in their abundance (or levels), as these do not inform on the underlying pathways driving these changes. Since the deposition (and removal) of RNA modifications is linked to the lifecycle of the RNAs themselves (38–40), the change in methylation levels can result from changes either in RNA (de)methylation rates and/or in the transcription or decay of methylated RNAs, From this lens, methods are needed that can assess transcriptional and post-transcriptional effects on methylated RNAs under non-stationary conditions.

Stable isotope labeling is a well-established method to quantify metabolic activity in cultured cells (41–43). Here, we developed a quantitative approach using [^13^C-methyl]-methionine labeling and mass spectrometry (MS) to assess the turnover of base modifications (^13^C-dynamods) in newly transcribed RNA. With ^13^C-dynamods, we trace the proportion of newly methylated ribonucleosides and their decay through time, i.e. methylation turnover, from digested polyadenylated RNA and ncRNA. We first showed that polyadenylated RNA and ncRNA were distinguished by the different turnover frequencies (in hr^-1^) of modified ribonucleosides, which are inherently linked to the different half-lives of mRNA, rRNA and tRNA (39, 44, 45). Examining the kinetics of methylation turnover at steady-state within and across RNA classes as well as in free ribonucleosides enabled us to resolve the origin of RNA modifications in digested RNA, and thereby uncover the presence of modifications in uncharacterized RNA classes, such as *N6,N6*-dimethyladenosine (m^6^_2_A). We then applied ^13^C-dynamods in conjugation with abundance measurements of modified ribonucleosides in polyA+ and ncRNA, which resolved their transcriptional versus posttranscriptional sensitivity in response to actinomyin D and metabolic stresses. Thus, the quantitative nature of ^13^C-dynamods demonstrates its capacity for sensitive characterization of modified ribonucleosides, their origin and dynamics, in multiple RNA classes of interest.

## RESULTS

### ^13^C labeling of polyadenylated and ribosomal RNA modifications

S-adenosylmethionine (SAM) is the direct substrate of RNA methylation reactions in eukaryotic cells (46). To trace the incorporation of SAM into RNA, we cultured 786O cells in methionine-free DMEM medium supplemented with either unlabeled methionine (‘Unlab’) or [^13^C-methyl]-methionine, and analyzed the isotopologues (m+0, m+1, m+2) of modified and unmodified ribonucleosides by tandem mass spectrometry (LC-MS/MS) (Fig. 1A, 1B). The m+0 isotopologue (e.g. 150.1 *m/z* for m^6^A) represents the mass of the analyzed molecule where all atoms are present as the most common isotope, whereas the m+1 isotopologue (e.g. 151.1 *m/z* for m^6^A) indicates the mass shift due to ^13^C incorporation from the ^13^C-labelled methionine tracer or from the natural abundance of ^13^C, ^15^N, ^18^O and ^2^H stable isotopes. With ^13^C-dynamods, we measure the isotopologue fractions [m+1 / (m+1 + m+0)] of modified and unmodified ribonucleosides, which reflect the amount of ^13^C enrichment (m+1) in the measured ribonucleoside relative to the total pool of that ribonucleoside (m+1 + m+0) (Fig. 1A). Thus, the isotopologue fractions of each ribonucleoside during [^13^C-methyl]-methionine labeling are internally controlled for the amount of pre-existing ribonucleosides (m+0 and naturally labelled m+1) prior to ^13^C labeling. Here, we analyzed the isotopologues of *N*6-methyladenosine (m^6^A), 7-methylguanosine (m^7^G), 1-methyladenosine (m^1^A), *N*6,*N*6-dimethyladenosine (m^6^ _2_A), 2’-O-methyladensone (A_m_), 5-methylcytosine (m^5^C) and unmodified ribonucleosides from digested polyadenylated (polyA+), large (>200 nt) and small (<200 nt) RNA. We observed increased m+1 and concomitant decreased m+0 ion counts in modified ribonucleosides from polyA+ and large RNA after 4 and 24 hours of [^13^C-methyl]-methionine labeling, while the m+1 fractions (natural abundance) of the unmodified ribonucleosides was unaltered during the labeling period (Fig. 1C, 1D; fig. S1A; Supplementary Materials). We determined that the ^13^C enrichment of intracellular methionine and SAM reaches a 98-100% plateau within 30 minutes and remains constant thereafter (fig. S1B). Thus, the change in the ‘heavy’ (m+1) isotopologue fraction relative to the unlabelled condition (natural abundance) indicates SAM-dependent RNA methylation of newly synthesized RNA, whose turnover we have examined within and across RNA classes. In contrast to singly methylated ribonucleosides, the m^6^_2_A modification exhibited an enrichment mostly of the m+2 isotopologue upon ^13^C labeling (Fig. S1A), and so the m+2 fraction was used to assess m^6^_2_A dynamics in subsequent analyses.

**Figure 1.**
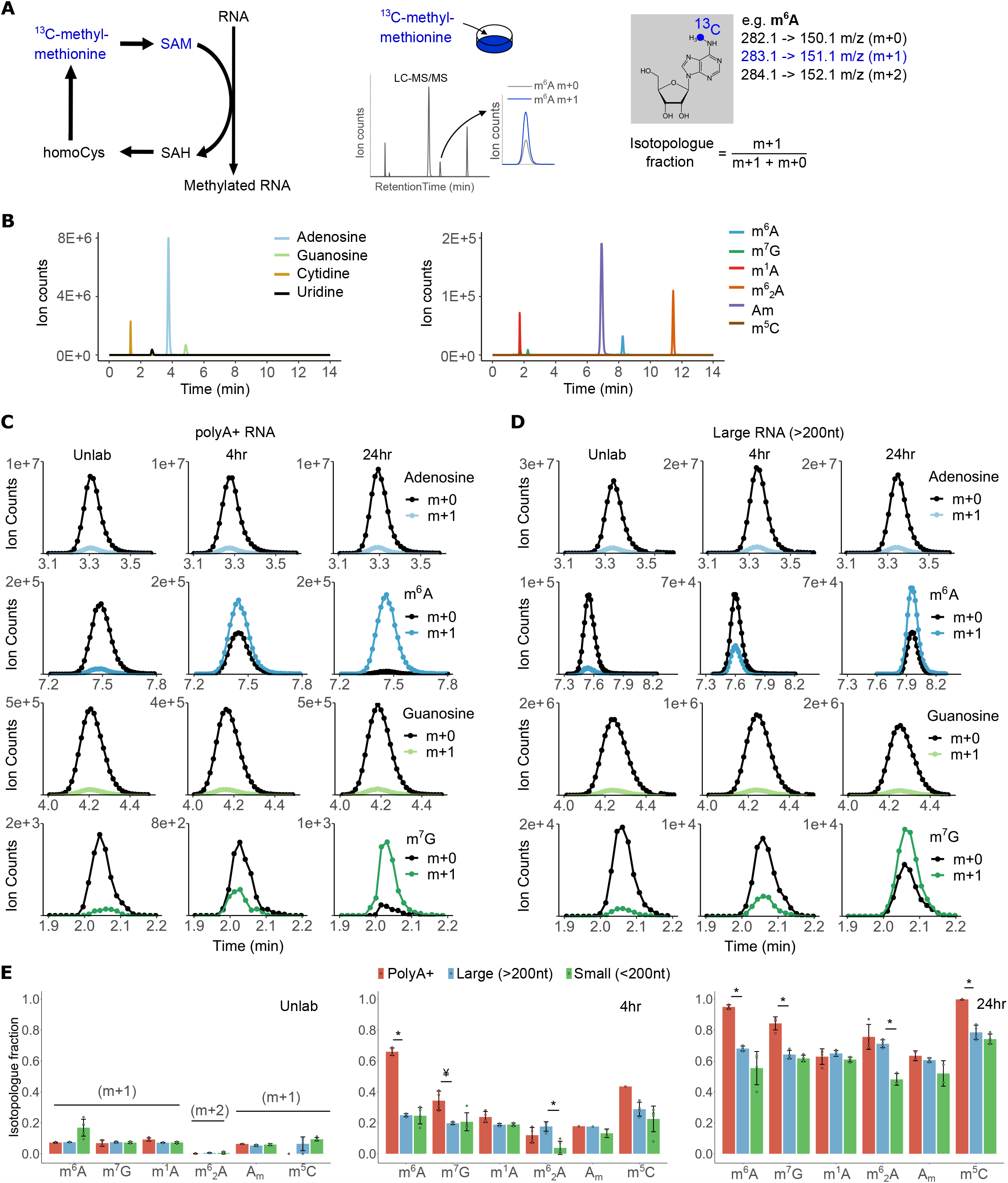
^13^C labelling of RNA modifications across RNA classes. (**A**) The ^13^C-dynamods workflow shows the tracing of RNA methylation: cells are cultured with [^13^C-methyl]-methionine, the RNA is isolated, digested to ribonucleosides and subjected to LC-MS/ MS analysis. The isotopologues detected for m^6^A are shown. (**B**) Representative chromatogram of unmodified and modified ribonucleosides from total RNA. (**C-D**) The m+0 and m+1 isotopologues of modified and unmodified ribonucleosides (representative chromatograms) in polyA+ (C) and large RNA (D). (**E**) Quantification of the isotopologue fractions of each ribonucleoside in polyA+, large and tRNA under unlabelled (‘Unlab’) conditions, after 4 and 24 hours of culture with [^13^C-methyl]-methionine. m^6^A, *N*6-methyladenosine; m^7^G, 7-methylguanosine; m^1^A, 1-methyladenosine; m^6^_2_A, *N*6,*N*6-dimethyladenosine; Am, 2*’*-O-methyladenosine; m^5^C, 5-methylcytidine; m^1^G, 1-methylguanosine; m^2^G, 2-methylguanosine. SAM, s-adenosylmethionine; SAH, s-adenosylhomocysteine; homoCys, homocysteine. Error bars represent standard deviation of three to four biological replicates, with the exception of m^5^C in polyA+-purified RNA (two replicates at 24hr, one replicate in the Unlab/4hr time points). In all cases, each replicate is the average of two technical replicas. * denotes P < 0.005, ¥ denotes P < 0.05, of a two-sided Student’s t-Test comparing samples as indicated.

Quantification of isotopologue fractions showed a faster kinetics for m^6^A, m^7^G and m^5^C in polyA+ RNAs when compared to large RNA (>200nt) and small RNAs (<200 nt) (Fig. 1E), in accordance with the faster lifecycle of mRNA in mammalian cells (45). In contrast, the kinetics of m^1^A, m^6^_2_A and Am methylation was similar between polyA+ and large RNA fractions (Fig. 1E), suggesting that a large portion of the signal for these modifications in the polyA+ fraction might derive from contaminating ribosomal RNA (rRNA). Of note, the isotopologues are analyzed on all ribonucleosides from the same polyA+ pool (Fig. S1C), so the purity of polyA+ enrichment is the same in all cases. This indicates that the relative abundance in mRNAs vs contaminating ncRNA is much higher for m^6^A, m^7^G and m^5^C compared to m^1^A, m^6^_2_A and Am. Interestingly, we detected m^6^_2_A in small RNAs from 786O cells, and its kinetics was different when compared to large RNA (Fig. 1E).

### Methylation turnover to resolve the origin of RNA modifications and their presence in uncharacterised RNA classes

To examine the kinetics of methylation turnover, we cultured cells for 24 hours with [^13^C-methyl]-methionine followed by replacement (chase) with naturally labelled methionine and analyzed the isotopologues over time. In accordance with the preceding findings (Fig. 1E), we observed an exponential decay of the m+1 isotopologue fraction for m^6^_2_A/m^7^G but not for Am/m^1^A/m^6^ A in polyA+ RNA (Fig. 2A). Conversely, the modifications of total/large RNAs exhibited uniformly slow turnover (Fig. 2B), consistent with the high stability of rRNA and tRNA in growing cells (44). To test the kinetic behavior underlying methylation turnover, we examined the goodness-of-fit of a linear versus exponential regression of the isotopologue fraction. Analysis of residual errors showed that the linear regression (m_(t)_ = m_(0)_ + kt) fits well the turnover of m^1^A/m^6^ A/A_m_ but not m^6^_2_A/m^7^G in polyA+ RNA, in which the ‘U-shaped’ curve supports a non-linear model (m_(t)_ = m_(0)_ e^(-kt)^) (Fig. 2C; fig. S2A). Conversely, the linear regression fitted well the turnover of ncRNA modifications (fig. S2B, S2C). The turnover frequency determined for Am/m^1^A/m^6^_2_A in polyA+ was similar to ncRNA modifications (k = 0.031 hr^-1^ on average), and significantly slower than m^6^A (k = 0.244 hr^-1^) or m^7^G (k = 0.089 hr^-1^) in polyA+ RNA (Fig. 2A, 2B). Thus, the turnover of m^1^A/m^6^_2_A/A_m_ from polyA+ fractions is incompatible with the exponential turnover of mRNA modifications (44, 45), confirming that they originate from ncRNA contamination.

**Figure 2.**
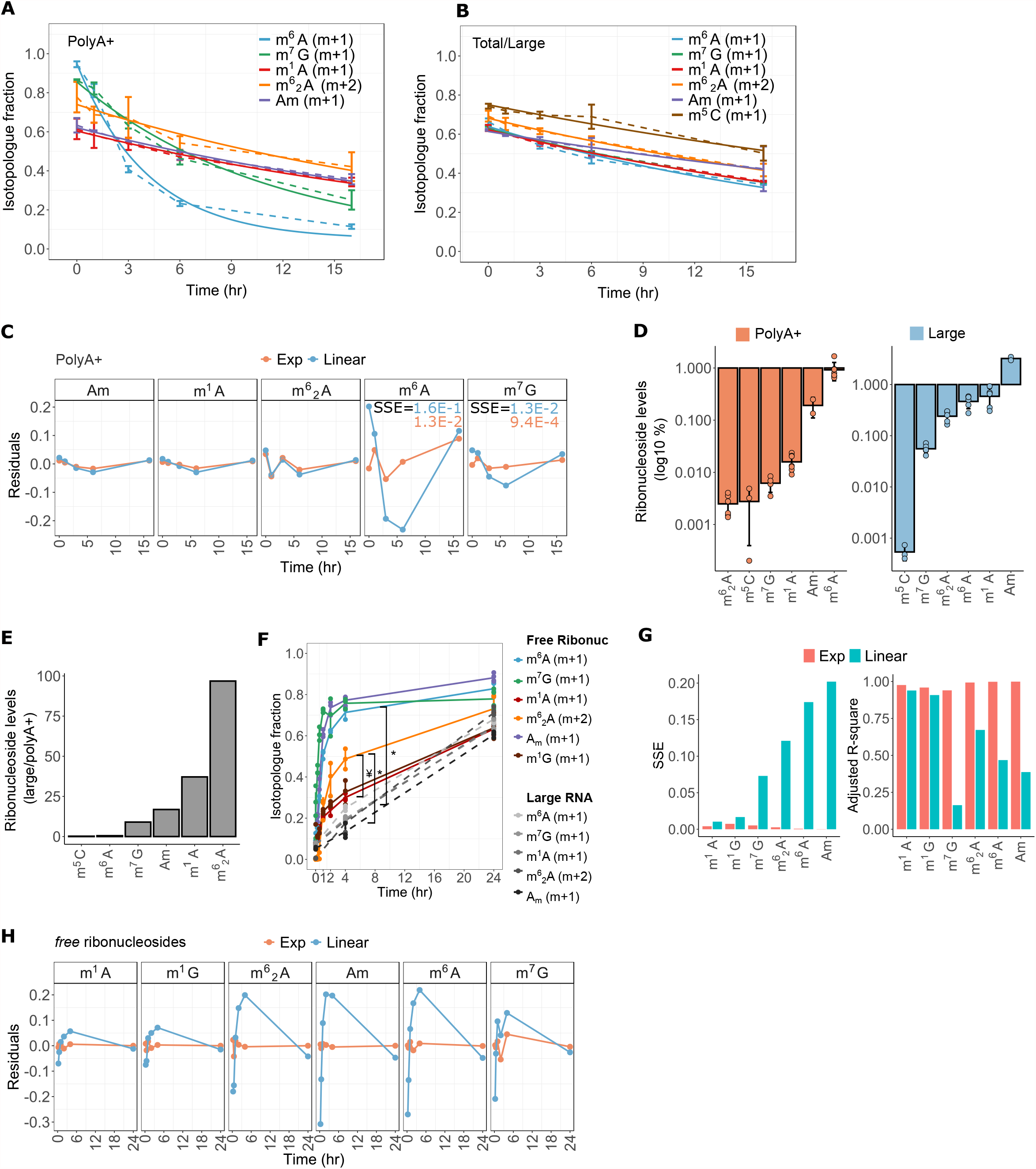
Turnover of modified ribonucleosides in polyA+, ncRNA and free ribonucleoside pool. (**A-B**) Isotopologue fractions during the ‘chase’ of ^13^C-labelled modifications with naturally labelled methionine for 0, 1, 3, 6 and 16 hours, in polyA+ (A) and total/ large RNA (B); dashed lines connect data points; solid lines, exponential (A) or linear (B) fit of isotopologue fractions. (**C**) Residuals of a linear vs. exponential regression of isotopologue fractions in polyA+ RNA in the chase experiment. (**D**) Normalized ion counts of modified ribonucleosides relative to the ion counts sum of all ribonucleosides, shown for polyA+ and large RNA. (**E**) Ratio of normalised ion counts between polyA+ and large RNA, as determined in (D). (**F**) Isotopologue fraction of free ribonucleosides analysed from metabolic extracts; grey datapoints refer to large RNA modifications (Fig. 1E) for comparison. (**G-H**) Goodness-of-fit of a linear vs. exponential fit of the isotopologue fractions in free modified ribonucleosides. SSE, sum of squared errors. m^6^A, *N*6-methyladenosine; m^7^G, 7-methylguanosine; m^1^A, 1-methyladenosine; m^6^_2_A, *N*6,*N*6-dimethyladenosine; A _m_, 2’*-O-*methyladenosine; m^5^C, 5-methylcytidine; m^1^G, 1-methylguanosine; A, adenosine; G, guanosine. Error bars represent 90% confidence intervals in (a-b), or standard deviation in (d, f) of at least three biological replicates with exception of A_m_ (two replicates). In all cases, each replicate is the average of two technical replicas. * denotes P < 0.005, ¥ denotes P < 0.05, of a two-sided Student’s t-Test comparing m^6^_2_A and m^6^A in the free ribonucleoside pool (free ribonuc) to large RNA, or m^6^2 A to m^1^A in the frree ribonuc pool.

The m^7^G modification was similar to less abundant than m^1^A/Am/m^6^_2_A in polyA+ RNA (Fig. 2D), and the sensitivity of MS detection was similar between m^7^G and m^1^A/Am/m^6^_2_A, as determined from equimolar injections of pure compounds (fig. S2D). Thus, the distinct turnover of RNA modifications in the polyA+-purified fraction was not explained by analytical sensitivity or normalised abundances alone. Instead, the modified ribonucleosides were more abundant in large RNA relative to polyA+ RNA by a factor of 17/37/97/9 for A_m_/m^1^A/m^6^_2_A/m^7^G, respectively, while m^6^A level was greater in polyA+ RNA by a factor of 2 (Fig. 2E). From the determined turnover frequencies (hr^-1^) of ncRNA (Fig. 2B), we estimated that a 50% contribution of ncRNA signal to the ribonucleoside pool in polyA+-purified RNA would lower the detected turnover of the bona fide m^6^A mRNA modification, with a turnover frequency of 0.242hr^-1^, to that of m^7^G, with a turnover frequency of ∼0.09 hr^-1^ (fig. S2E). That is, reliable turnover-based detection of a hypothetical mRNA modification that is more abundant by a factor of 17-97 in ncRNA relative to mRNA is attainable with a maximal ncRNA contamination of 0.5% (50%/97) to 2.9% (50%/17), but even an optimal performance of available methods to purify polyA+ RNA commonly contains 2-3% of contaminating ncRNA (fig. S2F) (36, 37). Thus, the ability to resolve bona fide mRNA modifications based on methylation turnover is limited by the depletion efficiency of highly abundant rRNA modifications.

As the turnover frequency of m^6^A was ∼8 times faster in polyA+ RNA than ncRNA (Fig 2A, 2B), we reasoned that free ribonucleosides derived from metabolic extracts would be derived mainly from endogenous degradation of mRNAs, rather than ncRNAs. Thus, we employed ^13^C-dynamods to examine the methylation turnover of free ribonucleosides, thereby enriching for mRNA-derived ribonucleosides and decreasing interference from rRNA-derived ribonucleosides. We found a fast kinetics following ^13^C labeling that was similar for m^6^A, m^7^G and A_m_, indicating that they are primarily derived from the same RNA type (i.e. mRNAs) (Fig. 2F, 2G, 2H). Interestingly, the m^6^_2_A modification also exhibited a non-linear, faster kinetics than m^1^A or 1-methylguanosine (m^1^G) (Fig. 2F, 2G, 2H), which are known tRNA modifications and were present in the metabolic extracts at high levels (fig. S2G) (47). We did not include the isotopologue analysis of m^5^C in the time-series experiments due to low abundance of its free ribonucleosides, and due to low MS sensitivity for m^5^C (Fig. 2D; fig. S2D, S2G), which compromises the quantification under conditions of partial ^13^C labeling. Thus, while the turnover analysis of the polyA+ fraction couldn’t be used to validate lowly abundant modifications due to rRNA contamination (Fig. 2E), the turnover of Am and m^6^_2_A in the free pool suggests that these modifications indeed partly derive from rapidly decaying, non-ribosomal RNAs (Fig 2F), which may include mRNAs. This was supported by the normalized levels of A_m_ and m^6^ _2_A in the free pool being more similar to those of polyA+ than rRNA (fig. S2H). These results confirm the presence of A_m_ in mRNA (48) and suggest m^6^_2_A is more common in short-lived RNAs than m^1^A, which has already been studied in mRNAs (49, 50).

### Sensitivity of RNA modifications to metabolic stresses

The maintenance of methylation reactions requires SAM and serine/glycine, which feed one-carbon metabolism and SAM synthesis (51). Conversely, RNA demethylation is catalysed by α-ketoglutarate (α-kg)-dependent dioxygenases such as FTO and ALKBH5, which act on m^6^A, *N*6,2*-O-*dimethyladenosine (m^6^A_m_) and m^1^A (52, 53). We measured both the methylation turnover and abundance of modified ribonucleosides to obtain insights into the sensitivity of mRNA versus ncRNA to metabolic stresses linked to RNA (de)methylation. First, we confirmed that actinomycin (ActD), a pan inhibitor of eukaryotic transcription, completely inhibited the methylation turnover of polyA+ and large RNA (Fig. 3A), in accordance with the co-transcriptional deposition of mRNA and most rRNA modifications (54–57). Treatment with ActD decreased the normalised levels of m^6^A (m^6^A/A) in polyA+ RNA (Fig. 3B), but not of m^7^G, which supports the role of m^6^A in global mRNA destabilisation, as initially reported (40). Deprivation of serine or glutamine inhibited mainly the methylation turnover of large RNA (Fig. 3C, 3D). In contrast, the abundance of modified ribonucleosides in large RNA was unaltered under these forms of stress (Fig. 3E) whereas glutamine deprivation led to increased m^6^A levels in polyA+ RNA (Fig. 3F), which was in contrast to ActD treatment (Fig. 3B). Thus, the reduced methylation turnover of large RNA simply results from an inhibited transcription of the rRNAs themselves, while the altered abundance of m^6^A without significant changes in its turnover indicates its post-transcriptional lability to stress conditions via altered i) decay of m^6^A-enriched vs. m^6^A-depleted mRNAs or ii) dynamics of the m^6^A modification itself. These data show that measurements of methylation turnover and ribonucleoside abundances resolve transcriptional versus post-transcriptional effects on RNA modifications in non-stationary (stimulus-dependent) experiments.

**Figure 3.**
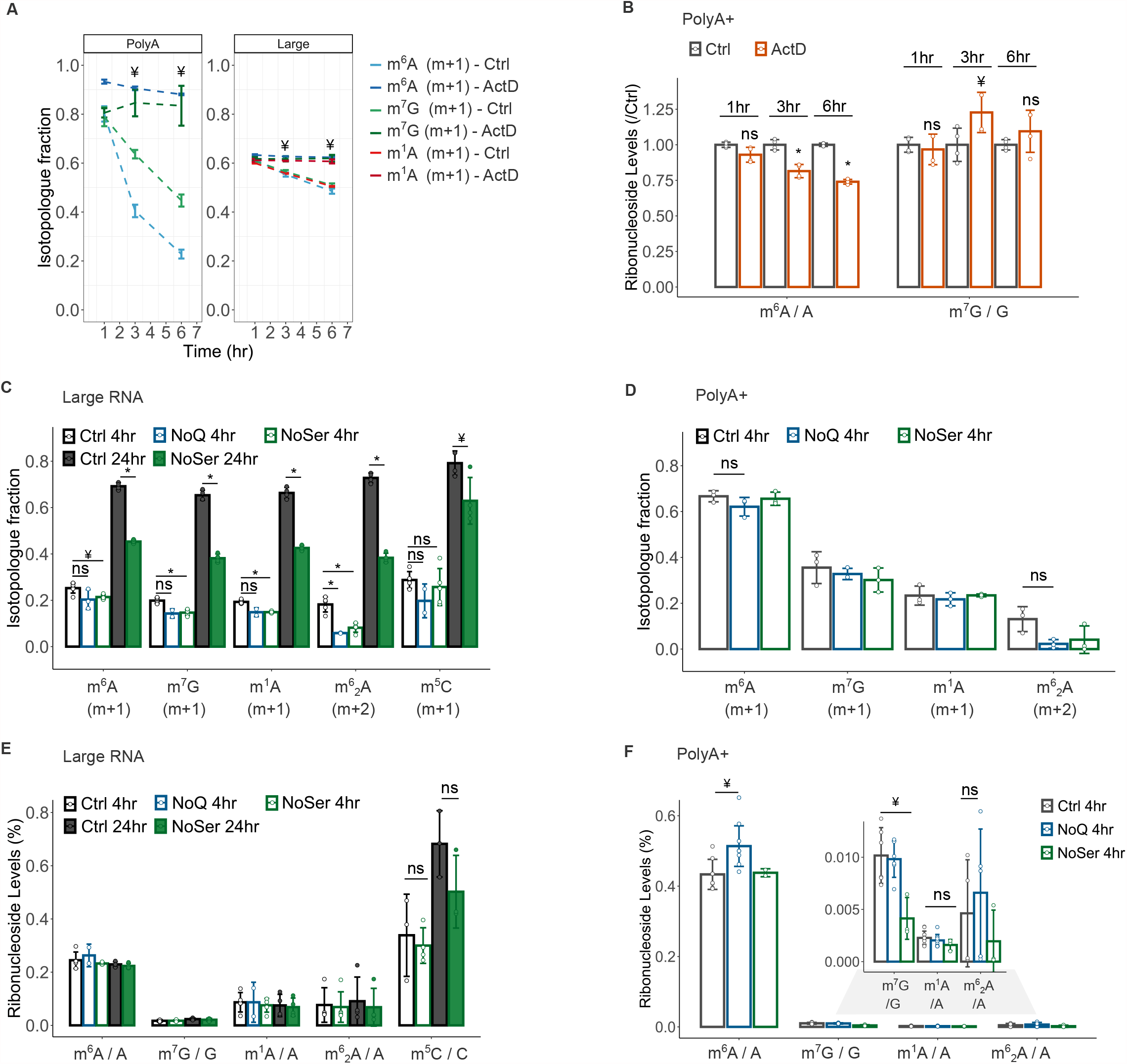
Dynamics of RNA modifications under stress conditions. (**A**) Isotopologue fractions of modified ribonucleosides in polyA+ and large RNA during a ^13^C-dynamods chase experiment in 786O cells cultured with complete DMEM medium (ctrl) or DMEM supplemented with actinomycin D (ActD, 10µM); dashed lines connect data points. (**B**) Normalised levels of m^6^A (m^6^A/A) and m^7^G (m^7^G/G) in polyA+ RNA derived from ctrl and ActD-treated 786O cells, normalised relative to the ctrl data points in the chase experiment as indicated. (**C-D**) Isotopologue fractions of modified ribonucleosides in large RNA (C) and polyA+ RNA (D) following [^13^C-methyl]-methionine labelling under control (ctrl), glutamine-(NoQ), or serine-(NoSer) deprived conditions. (**E-F**) Normalised levels of modified ribonucleosides in large RNA (E) and polyA+ RNA (F) in 786O cells cultured with ctrl, NoQ or NoSer DMEM medium for the times indicated. Error bars represent standard deviation of three or more biological replicates, with exception of (A-B), the NoQ condition in (C, E) and m^6^A under NoSer (D, F) (two biological replicates). In all cases, each replicate is the average of two technical replicas. * denotes P < 0.005, ¥ denotes P < 0.05, ns denotes not significant, of a two-sided Student’s t-Test comparing stress to ctrl samples, as indicated.

## DISCUSSION

Isotopic labeling of cultured cells is a well-established method to quantify metabolic activity, but its application to RNA modifications has been limited to ncRNAs (33, 58). Here, we demonstrate that quantification of methylation turnover with dynamic ^13^C labeling informs on the distinct dynamics of polyA+ and ncRNA modifications, and their sensitivity to metabolic perturbations of mammalian cells. ^13^C-dynamods presents several advances in the application of mass spectrometry (MS) approaches to study RNA turnover & dynamics. First, in contrast to approaches that rely on prior/in-parallel labeling with ^15^N/^13^C-enriched nucleosides to distinguish between pre-existing and newly modified RNA (33, 58), the ‘heavy’ isotopologue fraction derived from [^13^C-methyl]-methionine is internally controlled for the amount of unlabelled, pre-existing ribonucleosides, and is thus specific to modified ribonucleosides. Second, in contrast to approaches using multiplexing or spiking from isotopically labelled cultures (24, 25, 32, 58), the dynamic ^13^C labeling over time directly informs on the turnover frequency (in hr^-1^) (41, 43) of various RNA modifications, as the direct substrate (SAM) of the targeted reaction (RNA methylation) is close to 100% labelled in less than 30 minutes in cell culture (fig. S1B) (43). Moreover, SAM needs not be fully labelled for comparisons of dynamic ^13^C labeling between modifications at early, single time points as its ^13^C enrichment is expected to be equal across SAM-dependent RNA modifications. Third, conventional RNA labeling approaches using ribonucleoside analogues or precursors rely on their incorporation into newly synthesized RNA through the salvage pathway (e.g. uridine and its analogs) (45, 59) or the *de novo* pathway (glucose, serine/glycine or water tracers) of ribonucleotide (NTP) synthesis (60–64). In contrast, the methylation of newly transcribed RNA occurs downstream (co-transcriptionally) of NTP synthesis, making it a reliable readout of RNA turnover, particularly under non-stationary conditions that may alter the relative activity of NTP synthesis pathways (65, 66). Also, the nucleotide recycling through the salvage pathway may lead to ineffective ‘washout’ with unlabelled nucleotides in chase experiments (59, 66), whereas a new SAM molecule is used in each (co-transcriptional) methylation cycle in ^13^C-dynamods, adding to the specificity and versatility of experimental designs (59). Finally, combined measurements of methylation turnover and ribonucleoside levels allow ^13^C-dynamods to also inform on transcriptional versus posttranscriptional sensitivity of RNA modifications in response to a stimulus.

Reliable study of mRNA modifications by MS analyses of ribonucleosides is challenging due to the high abundance of heavily modified ncRNAs that invariably contaminate the polyA+ fraction (36, 37). To address this challenge, we exploited the fact that the lifecycle of methylated RNA follows the lifecycle of the RNAs at metabolic steady-state (38–40). The decay of mRNA is fast with a median half-life from 40 min to 9 hours in mammalian cells (44, 67, 68), while rRNA and tRNA exhibit half-lives of 60-70 hours in growing fibroblasts (44). By definition, the methylation turnover as determined by ^13^C incorporation into newly methylated RNA does not assess non-modified RNAs, and hence does not capture the removal of RNA modifications. That is, any RNA demethylation activity can only affect the abundances of modified ribonucleosides as a whole (sum of m+1 and m+1), not the m+1 or m+0 isotopologues individually, which would be required to cause a differential m+1 fraction. Since RNA transcription and decay rates are constant at metabolic steady-state, we propose that the methylation turnover is an accurate readout of RNA turnover, which allowed us to assess the contributions of short-lived (mRNAs) and long-lived (non-coding) RNAs to a pool of modified RNAs and free ribonucleosides. Indeed, this information could not be drawn if labelled ribonucleoside themselves or its precursors were used to trace ribonucleotide synthesis (via the salvage and *de novo* pathways) or transcription. Likewise, the source RNA cannot be pinpointed if the abundance (levels) of modified ribonucleosides measured in isolation.

Using this approach, we could not reliably determine highly abundant rRNA modifications (Am/m^1^A/m^6^ A) in polyA+-purified RNA as being derived from mRNAs. Based on methylation turnover, we estimated that these modifications in the polyA+ fraction could be detected with a maximal ncRNA contamination of 0.5-2.9%, since Am/m^1^A/m^6^_2_A levels were found higher in ncRNAs by a factor of 17-37-97, respectively. In contrast, a ∼6% of ncRNA contamination would be required to fully account for the slower m^7^G turnover detected in polyA+ RNA, since m^7^G levels were found only ∼9 times higher in ncRNAs. As the m^7^G turnover reflected that of a bona fide mRNA modification, its slower kinetics in polyA+-purified RNA suggests a temporal delay relative to m^6^_2_A turnover. A further evidence for the distinct turnover of m^7^G relative to m^6^A in mRNAs is the response to metabolic stresses, where the m^7^G turnover changed in large RNA but not polyA+ RNA following 4 hours of ^13^C labeling (Fig. 3c, 3d). In this respect, while m^6^A is co-transcriptionally deposited in mRNA (39, 54, 56), m^7^G is an essential modification at the 5*’* cap of mRNA that can be placed both in the nucleus and cytoplasm (69, 70). Moreover, m^7^G sites have been reported internally within mRNA, catalysed by the cytosolic METTL1 methyltransferase (71). Taken together, the distinct kinetics of m^7^G and m^6^A in polyadenylated RNA suggest compartmentalization differences affecting their temporal deposition into newly synthesized mRNA.

Previous reports identified m^5^C and Am as mRNA modifications (48, 72), and more recently also m^1^A (49, 50). Our analysis of free ribonucleosides showed a similarly fast kinetics for the turnover of Am, m^6^A and m^7^G, consistent with these modifications being derived from the rapidly decaying mRNAs. This was supported by the normalized levels of free modified ribonucleosides being more similar to those of polyA+-purified than of ncRNAs. In contrast, the turnover of free m^1^A and m^1^G ribonucleosides was significantly slower, indicating that these modifications are predominantly derived from a ncRNA class, likely tRNAs, where they are present at high levels (47). This aligns with the recent conclusions that m^1^A might be restricted to a handful of mRNAs (3, 49). Finally, despite the low sensitivity of m^5^C detection in chase experiments, the quantification of m^5^C isotopologues at 24 hours is consistent with it being derived from mRNA (72).

The m^6^_2_A modification is thought to be present primarily in the 18S and 12S rRNA of mammalian transcriptomes (26, 73, 74). While m^6^_2_A has been detected in bacterial tRNA (75), care must be taken with abundant m^6^_2_A-modified rRNA fragments that co-purify with tRNA (76), and are not resolved by standard MS quantifications. Here, we detected slower turnover of m^6^_2_A in mammalian small RNAs relative to large RNA, raising the possibility that this modification is present in non-18S/12S RNA species, e.g. in small rRNA and/or tRNAs. Nevertheless, an unlikely possibility from ^13^C-dynamods measurements remains that m^6^_2_A-containing 18S/12S rRNA fragments are turned over at a slower rate than m^6^_2_A-containing intact rRNAs. Interestingly, the turnover of free m^6^_2_ A ribonucleosides exhibited a faster kinetics than free m^1^A/m^1^G (canonical ncRNA modifications) (47) and when compared to large RNA modifications, suggesting that m^6^_2_A is also present in RNAs of high turnover RNAs, which likely include mRNAs and short-lived ncRNAs. The presence and role of m^6^_2_ A beyond the 18S/12S rRNAs thus merits further investigation. Of note, single methylation intermediates of m^6^_2_ A have been detected *in vitro* (77), but we did not detect m+1 isotopologues for m^6^ A above its natural abundance. As SAM was 98-100% ^13^C-labelled within 30 minutes, the time resolution of our experiments does not capture sequential methylation of adenosine into m^6^_2_A. These various findings highlight the value examining the turnover of methylated RNA to uncover the presence of modifications in uncharacterised in RNA subclasses, which warrants further investigation.

A non-stationary condition, e.g. a stress condition, may affect the afferent (i.e. methylation of newly transcribed RNA) or efferent (i.e. decay of methylated RNAs) turnover parts of methylated RNAs. Combined with abundance measurements, this allowed us to assess transcriptional versus non-transcriptional sensitivity of modified RNAs to stress conditions. This was evidenced by the reduced methylation turnover of ncRNAs following serine deprivation and ActD treatment, wherein the abundance of modifications was unaltered in the former (transcriptional effect) but m^6^A levels decreased in the latter (posttranscriptional effect). The observed inhibited turnover of methylated ncRNAs under serine deprivation is in line with the expected inhibition of mTOR activity, and thereby of rRNA biogenesis (78). In contrast to ncRNAs, glutamine deprivation increased m^6^A levels in polyA+-purified RNA without significantly changing (not increasing) its turnover. These data suggest that m^6^A is post-transcriptionally sensitive to glutamine levels through either i) an altered decay of m^6^A-enriched vs. m^6^A-depleted RNAs, or ii) inhibition of m^6^A demethylation itself. The demethylation of m^6^A is a step-wise conversion into adenosine through the formation of *N*6-hydroxymethyladenosine (hm^6^A) and *N*6-formyladenosine (f^6^A) intermediates (79), whose presence or turnover was not assessed.

Thus, related developments are needed to address if post-transcriptional dynamics of m^6^A by metabolic stress conditions are due to RNA demethylation or differential decay of m^6^A-modified versus m^6^A-depleted unmodified RNAs. As glutamine is the main carbon source of α-ketoglutarate, a co-substrate of RNA demethylases (52, 53), it is plausible that its depletion could inhibit RNA demethylation. This supports the notion that the reversibility of m^6^A in mRNA is likely context-dependent (80). With exception of ActD treatment, the stimuli examined did not lead to changes in both the turnover and abundances of modified ribonucleosides. In this respect, while altered methylation turnover (and unaltered abundance) indicates a transcriptional effect, it is plausible that a stimulus may alter methylation turnover at the post-transcriptional level e.g. if the newly methylated (m+1 fraction) and pre-existing methylated (m+0 fraction) RNAs differently are exposed differently to RNA degradation, e.g. through nuclear/cytosolic compartmentalisation.

Our study demonstrates how quantification of methylation turnover and abundance can be used to examine the presence of RNA modifications in RNA classes, their temporal dynamics and sensitivity to metabolic stress conditions. These insights open new directions to be further explored by MS and orthogonal approaches to obtain information on particular RNAs.

## AUTHOR CONTRIBUTIONS

Paulo A. Gameiro: Conceptualization, Methodology, Formal analysis, Investigation, Resources, Data curation, Original Draft, Writing – Original Draft, Writing – Review & Editing, Visualization, Funding acquisition; Vesela Encheva: Methodology, Formal analysis; Mariana Silva Dos Santos: Formal analysis; James I MacRae: Resources, Writing – Review & Editing, Funding acquisition; Jernej Ule: Resources, Writing – Original Draft, Writing – Review & Editing, Supervision, Funding acquisition.

## ACKNOWLEDGMENTS

This project has received funding from Wellcome Trust (103760/Z/14/Z) and the European Research Council (ERC) under the European Union’s Seventh Framework Program (FP7/2007-2013)/ERC grant agreement ID: 617837). P.A.G was supported by a Marie Sklodowska-Curie individual fellowship (grant agreement ID: 701730). The Francis Crick Institute receives its core funding from Cancer Research UK (grant no. FC001002), the UK Medical Research Council (grant no. FC001002) and the Wellcome Trust (grant no. FC001002).

## CONFLICTS OF INTEREST

The authors declare that they have no competing interests.

## EXPERIMENTAL PROCEDURES

### Cell culture and metabolic labeling

786O cells were obtained from the Crick Cell Services and cultured at 37°C with 5% CO_2_ in high glucose DMEM medium (ThermoFisher Scientific, #61965026) supplemented with 10% fetal bovine serum (FBS) (ThermoFisher Scientific, #21875034). For ^13^C labeling experiments, 786O cells were grown in high glucose DMEM medium without glutamine, methionine and cystine (ThermoFisher Scientific, #21013024) supplemented with 10% dialyzed FBS (ThermoFisher Scientific, #26400044), 2 mM glutamine, 0.1 mM cystine and 0.2 mM [^13^C-methyl]-methionine (CK Isotopes Limited). Cells were maintained at 50-60% confluence (or 30-40% in chase experiments) and washed with PBS before switching to ^13^C-labelled medium (or regular medium in chase) for the indicated time periods.

### RNA purification

At the conclusion of metabolic labeling, the medium was aspirated and extraction of total RNA was performed with the mirVana isolation kit according to manufacturer’s instructions (Thermofisher Scientific, #AM1560). Large (>200nt) RNAs were purified by adding 1/3 volume of 100% ethanol to the aqueous phase recovered from the organic extraction before loading into the filter cartridge of the mirVana Kit. Small (<200nt) RNAs were purified by collecting the total filtrate, addition of 2/3 volume of 100% ethanol and loading into the filter cartridge. Polyadenylated RNA was purified from total/large RNA via two rounds of polyA tail hybridization with Oligo-dT magnetic Dynabeads (ThermoFisher Scientific, #61002).

### LC-MS/MS analysis of ribonucleosides

Purified RNA (100-250 ng) was digested into ribonucleosides using one unit of nuclease P1 (Sigma-Aldrich, #N8630-1VL) in 25 μl of buffer 25 mM NaCl, 2.5 mM ZnCl_2_ and 10mM NaCH_3_COO pH 5.3 and incubated for 2 hours at 37°C. Subsequently, NH_4_HCO_3_ (100 mM) and 5 units of alkaline phosphatase (CIP) (NEB, #M0525S) were added and the sample incubated for 2 hours (or 20 min, with Quick CIP) at 37°C. Formic acid was added at 0.1% v/v in a final volume of 50 μl and samples were filtered (0.22 μm, Millipore) and 15-20 μl analyzed in duplicate by LC-MS. Ribonucleosides were resolved with a C18 reverse phase column (100 × 2.1 mm, 3 μm particle size, Chromex Scientific, #F18-020503) and eluted with a gradient of 0.1% v/v formic acid (solvent A) and 80% acetonitrile in 0.1% formic acid (solvent B) at a flow rate of 0.2 ml/min and 40°C: 100% solvent A for 3 min, 12% solvent B for 12 min, and 100% solvent B for 2 min after which the column was re-equilibrated with 100% solvent A for 3 min (20 min total run time). The ribonucleoside separation was performed using U3000 HPLC (Thermo Scientific) and the detection by a TSQ Quantiva Triple Quadrupole mass spectrometer (TSQ Quantiva, Thermo Scientific) controlled by the Xcalibur software version 4.0.27 (Thermo Scientific). The HPLC was coupled to the TSQ Quantiva using a HESI (heated electrospray) ion source (Thermo Scientific) operating in positive ionization mode with the following parameters: capillary voltage, 3500V; sheath gas flow, 7.35 l/min; gas temperature, 325C. The first and third quadrupoles (Q1 and Q3) were stringently fixed to 0.2 units of resolution and set to detect the mass of the precursor ribonucleoside ion (Q1) and of the base and ribose product ions (Q3). The ribonucleosides were identified by comparison of the retention time and detected mass transitions to commercially available standards. The collision energies were experimentally defined based on the fragmentation pattern of each ribonucleoside standard and chosen based on the maximum intensity of the base product; the ribose ring was used only as a qualifying transition. The retention time, mass transitions (*m/z*) and collision energies of each ribonucleoside were: adenosine, ∼4.1 min, 268.1 -> 136.1 *m/z*, 20 V; guanosine, ∼5.4 min, 284.1 -> 135.0 *m/z*, 35.5 V; cytidine, ∼1.4 min, 244.2 -> 112.05 *m/z*, 12 V; m^6^A, ∼8.7 min, 282.1 -> 150.1 *m/z*, 20 V; m^1^A, ∼1.8 min, 282.1 -> 150.1 *m/z*, 20V, m^7^G, ∼2.3 min, 298.05 -> 166.1 *m/z*, 20V; m^5^C, ∼1.8 min, 258.2 -> 126.1, *m/z*, 13V; m^6^ _2_ A, ∼11.9 min, 296.2 -> 164.1 *m/z*, 22V. Each mass transition above corresponded to the m+0 isotopologue, and increased by one (m+1), two (m+2) and three (m+3) units for detection of the other isotopologues *e*.*g*. m^6^ _2_A: 297.2 -> 165.1 (m+1), 298.2 -> 166.1 (m+2), 299.2 -> 167.1 (m+3). The dwell time for each transition was 30 ms for a duty cycle of 930 ms (31 transitions), and 8 to 20 data points per chromatographic peak were obtained for ‘short’ and ‘long’ peaks, respectively. A mix of ribonucleoside standards containing 0.5, 1, 5, 10, 50, 100, 500 fmol, 1, 5, 10, 50 or 100 pmol of each ribonucleoside was run in parallel after the biological samples for absolute quantifications, a subset of which is shown in Extended Data Fig. 2d. Data were recorded using the Xcalibur 3.0.63 software (ThermoFisher Scientific) and analyzed using Skyline (version 19.1) (81) (Supplementary Materials).

### Metabolite extraction and LC-MS analysis of SAM/free ribonucleosides

At the end of cell culture with [^13^C-methyl]-methionine, metabolic activity quenched by adding ice-cold PBS. Metabolites were extracted by addition of 600 μl ice-cold 1:1 (vol/vol) methanol/water to the cell pellets, samples were transferred to a chilled microcentrifuge tube containing 300μl chloroform and 600μl methanol (1500 μl total, in 3:1:1 vol/vol methanol/water/chloroform). Samples were sonicated in a water bath for 8 min at 4°C, and centrifuged (13000 rpm) for 10 min at 4°C. The supernatant containing the extract was transferred to a new tube for evaporation in a speed-vacuum centrifuge, resuspended in 3:3:1 (vol/vol/vol) methanol/water/chloroform (350μl total) to phase separate polar metabolites (upper aqueous phase) from non-polar metabolites (lower organic phase), and centrifuged. The aqueous phase was transferred to a new tube for evaporation in a speed-vacuum centrifuge, and resuspended in 100μl water for LC-MS acquisition. LC-MS analysis was performed using a Dionex UltiMate LC system (ThermoFisher Scientific) with a ZIC-pHILIC column (150 mm x 4.6 mm, 5 μm particle, Merck Sequant), as described previously (82). A 15 min elution gradient of 80% Solvent A (20 mM ammonium carbonate in Optima HPLC grade water, Sigma Aldrich) to 20% Solvent B (acetonitrile Optima HPLC grade, Sigma Aldrich) was used, followed by a 5 min wash of 95:5 Solvent A to Solvent B and 5 min re-equilibration. Other parameters were as follows: flow rate, 300 μL/min; column temperature, 25°C; injection volume, 10 μL; autosampler temperature, 4°C. All metabolites were detected across a mass range of 70-1050 *m/z* using a Q Exactive Orbitrap instrument (ThermoFisher Scientific) with heated electrospray ionization and polarity switching mode at a resolution of 70,000 (at 200 *m/z*). MS parameters were as follows: spray voltage 3.5 kV for positive mode and 3.2 kV for negative mode; probe temperature, 320°C; sheath gas, 30 arbitrary units; auxiliary gas, 5 arbitrary units. Parallel reaction monitoring (PRM) was used at a resolution of 17,500 to confirm the identification of metabolites; collision energies were set individually in HCD (high-energy collisional dissociation) mode. Data were recorded using the Xcalibur 3.0.63 software and analyzed using Tracefinder 4.1 (ThermoFisher Scientific) according to the manufacturer’s workflows.

### Quantification of methylation turnover

The isotopologue fractions were defined as the total ion counts of the m+1 isotopologue (except for m+2 in m^6^ _2_A) relative to the total ion counts of the m+0 plus m+1 isotopologues. The kinetics of isotopologue fractions and goodness-of-fit were determined using the Curve Fitting toolbox of Matlab R2020a (MathWorks) either by a linear regression [f(x) = p1*x + p2] or an exponential fit: a one-term function in the chase experiments [f(x) = a*exp(b*x), Levenberg-Marquardt algorithm] and a two-term function to fit the isotopologue fractions of free ribonucleosides [f(x) = a*exp(b*x) + c*exp(d*x), Levenberg-Marquardt algorithm].

## Notes

https://www.researchsquare.com/article/rs-109009/v2

### Competing Interest Statement

The authors have declared no competing interest.

